## Supplementary Text for "Distributed representations of prediction error signals across the cortical hierarchy are synergistic"

### Details of the neurocomputational model

### Microstructure

Each area consists of two neuronal layers, each of 625 (25x25) cells, one containing excitatory cells and one containing inhibitory ones (in what follows, referred to as e- and i-cells, respectively). To avoid any potential edge effects, layers have a toroidal structure: the top edge is adjacent to the bottom one, and the left edge is adjacent to the right one. In line with Wilson-Cowan models (Wilson & Cowan, 1973), a single pair of e- and i-cell models the average activity of a local population of pyramidal neurons and underlying inhibitory interneurons within one cortical column (grey matter under approximately 0.25 square mm of the cortical surface). Cells are modelled as graded-response neurons (see below).

Each e-cell is restricted to send projections to the 19x19 e-cell neighbourhood within the same area, to topographically corresponding 19x19 e-cell patches in connected areas, and to a 5x5 i-cell patch in the inhibitory layer of the same area (Fig. 1E). The probability of a synapse to be created between an e-cell and another cell falls off with their distance (Braitenberg & Schüz, 1998) according to a Gaussian function clipped to 0 outside the relevant neighbourhood. This produces a sparse, patchy and topographic connectivity, as typically found in the mammalian cortex (Amir et al., 1993; Kaas, 1997).

### Membrane dynamics

The state of an (excitatory or inhibitory) cell $e$ at time $t$ is uniquely defined by its membrane potential $V(e,t)$, determined by the following equation:

$$\tau \frac{dV(e,t)}{dt} = -V(e,t) + k_1(V_{in}(e,t) + k_2\eta(e,t)) \qquad (1)$$

where $V_{in}(e,t)$ is the sum of all postsynaptic inputs acting upon cell $e$ (see Eq. (2)), $\eta(e,t)$ is a white noise process with uniform distribution over [–0.5, 0.5], $\tau$ is the cell's membrane time constant (note that e- and i-cells have different $\tau$, see Table 1), and $k_1$ and $k_2$ are scaling constants. Note that the activity of each e-cell is intrinsically noisy, simulating the spontaneous baseline firing of real neurons (i-cells have $k_2$=0). The total input to a cell $e$ is defined as:

$$V_{in}(e,t) = (\sum E/IPSPs) - k_G\omega_G(e,t) \qquad (2)$$

where $\Sigma E/IPSPs$ is the sum of all excitatory and inhibitory postsynaptic potentials – I/EPSPs; inhibitory synapses are given a negative sign – acting upon neural cluster (cell) $e$ at time $t$, $\omega_G(e,t)$ is the global (or area-specific) inhibition (see Eq. (3)) and $k_G$ is a scaling constant. Note that each e-cell gets exactly one IPSP from its twin i-cell (see Fig. 1E).

The global inhibition mechanism is an area-specific inhibitory loop that prevents overall network activity from falling into non-physiological states (Braitenberg & Schüz, 1998). (Note that $k_G$=0 for i-cells: for simplicity, global inhibition acts only on e-cells). For each model area $A$, the global inhibition $\omega_G(e,t)$ is defined by:

$$\tau_G \frac{d\omega_G(e,t)}{dt} = -\omega_G(e,t) + \sum_{e \in A} O(e,t) \qquad (3)$$

where $\sum_{e \in A} O(e,t)$ is the sum of all e-cell outputs within area $A$ (see Eq. (4)) and $\tau_G$ is the global inhibitory response time constant.

All cells produce a graded response representing the average firing rate of the neural cluster; in particular, the output (transformation function) of an e-cell $e$ at time $t$ is defined as:

$$
O(e,t) = \begin{cases} 0 & \textit{if } V(e,t) \leq \varphi(e,t) \\ V(e,t) - \varphi(e,t) & \textit{if } 0 < (V(e,t) - \varphi(e,t)) \leq 1 \\ 1 & \textit{otherwise} \end{cases} \qquad (4)
$$

Eq. (4) above is a piecewise-linear sigmoid function of the e-cell's membrane potential $V(e,t)$, clipped into the range [0, 1] and with slope 1 between the lower and upper thresholds $\varphi(e,t)$ and $\varphi(e,t)$+1. The output $O(i,t)$ of an i-cell $i$ is 0 if $V(i,t) < 0$, and $V(i,t)$ otherwise (i.e., unlike e-cells, i-cells do not saturate, reflecting that real interneurons show little firing rate adaptation).

The threshold $\varphi(e,t)$ of an e-cell is not constant but depends on the cell's recent activity, so that the more active the cell, the higher the threshold (see Eq. (5)). This implements a simple form of homeostatic adaptation, or neuronal fatigue (Matthews, 2001):

$$
\varphi(e,t) = \alpha\, \omega(e,t) \qquad (5)
$$

where $\omega(e,t)$ is the estimated time-average of cell *e*'s recent output (see Eq. (6)) and α is a scaling constant (adaptation strength). The estimated time-average $\omega(e,t)$ of a cell's output is computed by integrating the following differential equation (Eq. (6)) with time constant $\tau_A$, assuming $\omega(e,t)$=0 at time $t$=0:

$$
\tau_A \frac{d\omega(e,t)}{dt} = -\omega(e,t) + O(e,t) \qquad (6)
$$

**Table S1** *Model parameters*

| | |
|---|---|
| $\tau_e = 2.5$ | e-cells membrane potential time constant (Eq. (1)) |
| $\tau_i = 5$ | i-cells membrane potential time constant (Eq. (1)) |
| $k_1 = 0.01$ | scaling constant (Eq. (1)) |
| $k_2 = 150\sqrt{(24/\Delta t)}$ | noise amplitude (Eq. (1)) |
| $\Delta t = 0.1$ | simulation step size (ms) |
| $\eta \sim U[-0.5, 0.5]$ | noise distribution (Eq. (1)) |
| $\tau_G = 60$ | global inhibition time constant (Eq. (3)) |
| $k_G = 95$ | global inhibition strength (Eq. (2)) |
| $\alpha = 100$ | adaptation strength (Eq. (5)) |
| $\tau_A = 50$ | e-cells estimated time-averaged activity time constant (Eq. (6)) |
